# Robustness of RADseq for evolutionary network reconstruction from gene trees

**DOI:** 10.1101/414243

**Authors:** José Luis Blanco-Pastor, Yann J.K. Bertrand, Isabel María Liberal, Yanling Wei, E.Charles Brummer, Bernard E. Pfeil

## Abstract

Although hybridization has played an important role in the evolution of many species, phylogenetic reconstructions that include hybridizing lineages have been historically constrained by the available models and data. Recently, the combined development of high-throughput sequencing and evolutionary network models offer new opportunities for phylogenetic inference under complex patterns of hybridization in the context of incomplete lineage sorting. Restriction site associated DNA sequencing (RADseq) has been a popular sequencing technique for evolutionary reconstructions of close relatives in the Next Generation Sequencing (NGS) era. However, the utility of RADseq data for the reconstruction of complex evolutionary networks has not been thoroughly discussed. Here, we used new molecular data collected from diploid perennial *Medicago* species using single-digest RADseq to reconstruct evolutionary networks from gene trees, an approach that is computationally tractable with datasets that include several species and complex patterns of hybridization. Our analyses revealed that complex network reconstructions from RADseq-derived gene trees were not robust under variations of the assembly parameters and filters. Filters to exclusively select loci with high phylogenetic information created datasets that retrieved the most anomalous topologies. Conversely, alternative clustering thresholds or filters on the number of samples *per* locus affected the level of missing data but had a lower impact on networks. When most anomalous networks were discarded, all remaining network analyses consistently supported a hybrid origin for *M. carstiensis* and *M. cretacea*.

## 1. Introduction

The reconstruction of a reticulate history in evolutionary close relatives has been considered from three different analytical perspectives: i) population genetic models including: approximate Bayesian Computation (Beaumont et al., 2002), full-likelihood genealogical samplers that make use of DNA sequences (Gronau et al., 2011; Hey, 2010; Sethuraman and Hey, 2016) and likelihood or pseudo-likelihood methods based on the joint allele frequency spectrum (Excoffier et al., 2013; Gutenkunst et al., 2009; Pickrell and Pritchard, 2012); ii) *D*- statistics (Durand et al., 2011; Eaton and Ree, 2013; Green et al., 2010; Meyer et al., 2012; Pease and Hahn, 2015); and iii) evolutionary network models (Solís-Lemus and Ané, 2016; Wen and Nakhleh, 2016; Yu et al., 2014, 2013; Yu and Nakhleh, 2015; Zhang et al., 2018; Zhu et al., 2017). The first two perspectives assume a previously known backbone phylogeny to formulate a hypothesis of hybridization. This backbone tree is usually constructed either using i) a total evidence approach with concatenation of full sequence information (Eaton and Ree, 2013; Escudero et al., 2014; Fernández-Mazuecos et al., 2017; Hipp et al., 2014; Wagner et al., 2013) or ii) coalescent based methods (Eaton and Ree, 2013; Fernández-Mazuecos et al., 2017; Rheindt et al., 2014) that reconcile individual gene trees. Despite being a standard approach, the construction of a backbone tree could be an incorrect representation of the main evolutionary history of the species under complex reticulate evolution (Clark and Messer, 2015; Huson et al., 2010; Yu et al., 2011), or molecular data can show a different “main” phylogeny when hybridization is first accounted for (Sousa et al., 2017).

Evolutionary networks provide an explicit model of evolutionary relationships that extends the tree model to allow for reticulations with internal nodes representing ancestral species. Recently developed phylogenetic network reconstruction methods are based on maximum parsimony (Yu et al., 2013), maximum likelihood (ML) (Yu et al., 2014), maximum pseudo-likelihood (Solís-Lemus and Ané, 2016; Yu and Nakhleh, 2015) and Bayesian inference (BI) methods (Wen et al., 2016; Wen and Nakhleh, 2016; Zhang et al., 2018; Zhu et al., 2017). Although ML and BI methods show promise, they are still limited to small datasets (usually fewer than 10 individuals and less than 3 reticulations, Yu and Nakhleh, 2015). In contrast, maximum pseudo-likelihood (summary) methods (Solís-Lemus and Ané, 2016; Yu and Nakhleh, 2015) are nowadays a convenient alternative for complex empirical datasets (Wen et al., 2017).

RADseq approaches (reviewed in Andrews et al., 2016) are widely used sequencing techniques for evolutionary reconstructions in the Next Generation Sequencing era. RADseq was first envisioned as a technique to find intraspecific genetic variation (Baird et al., 2008; Elshire et al., 2011; Hohenlohe et al., 2011). Later RADseq methods have been considered suitable for phylogenetic studies from shallow to deep timescales (Cariou et al., 2013; Eaton, 2014; Harvey et al., 2016; Rubin et al., 2012). RADseq are particularly appealing for systematics because they are easily applied to non-model organisms for which no reference genome or previous genomic information is available (Cariou et al., 2013; Fernández-Mazuecos et al., 2017; Rubin et al., 2012). For that reason RADseq has become a very popular technique for hybridization studies across a diversity of organisms and timescales (Escudero et al., 2014; Fernández-Mazuecos et al., 2017). Nevertheless, because RADseq datasets are limited in sequence length, contain relatively few variable sites, and do not generally yield resolved gene trees (Rubin et al., 2012), it is unknown if they are appropriate for maximum pseudo-likelihood phylogenetic network reconstruction methods. Their intrinsic characteristics suggest that these datasets are limited for network inference from gene trees, but an in-depth evaluation of their utility is still lacking.

*Medicago* L. (Fabaceae) is a genus comprising 87 species (Small, 2011) and includes the economically important forage crop alfalfa (*M. sativa*, section *Medicago*) in addition to the model legume *M. truncatula* (Barker et al., 1990; Benedito et al., 2008; Branca et al., 2011; Cook, 1999; Young et al., 2011). The genus *Medicago* L. exhibits severe phylogenetic gene tree incongruence that has been mainly attributed to hybridization and ILS (Eriksson et al., 2018, 2017; Maureira-Butler et al., 2008; Sousa et al., 2017, 2014; Steele et al., 2010; Yoder et al., 2013). We collected new molecular data from diploid perennial *Medicago* species using single-digest RADseq (Genotyping-By-Sequencing, Elshire et al., 2011). We investigated the ability of RADseq data to unveil the evolutionary history of diploid species of *Medicago* section *Medicago* using a network reconstruction method that uses gene trees (Yu and Nakhleh, 2015). Specifically, we investigated the robustness of this method (i.e. the propensity to retrieve a set of optimal networks with similar topologies) under a variety of RADseq data assembly parameters and filters.

## 2. Materials and Methods

### 2.1 Sampling

Our choice of species (Table 1) was based on results of previous studies grouping diploid perennial *Medicago* taxa (Bena, 2001; Maureira-Butler et al., 2008; Sousa, 2015; Yoder et al., 2013). These includes: *M. marina, M. cretacea, M. rhodopea, M. prostrata, M. daghestanica and M. sativa* (section *Medicago* subsection *Medicago*), *M. hybrida and M. suffruticosa* (section *Medicago* subsection *Suffruticosae*); *M. carstiensis* (section *Carstiensae*); *M. rugosa and M. scutellata* (section *Spirocarpos* subsection *Rotatae*). As outgroup we used the annual species *M. truncatula* (section *Spirocarpos* subsection *Pachyspirae*).

**Table 1.** Information on the *Medicago* samples used in the present study.

| Species | Accession No | Germplasm Bank | Country of origin | Accession European Nucleotide Database | Chromosome number<sup>d</sup> |
| --- | --- | --- | --- | --- | --- |
| *M. carstiensis* | PI641414 | USDA | Russian Federation |  | 16<sup>a</sup> |
| *M. carstiensis* | MED152/91 | Ösnabruck Botanic Garden | NA |  | 16<sup>a</sup> |
| *M. cretacea* | PI631721 | USDA | Russian Federation |  | 16<sup>b</sup> |
| *M. daghestanica* | Eoo337403; 1653*1 | GB herbarium | Russian Federation |  | 16<sup>a</sup> |
| *M. daghestanica* | Eoo337402; 1653*2 | GB herbarium | Russian Federation |  | 16<sup>a</sup> |
| *M. hybrida* | PI538998 | USDA | Russian Federation |  | <sup>d</sup> |
| *M. marina* | PI516711 | USDA | Morocco |  | 16<sup>a</sup> |
| *M. marina* | PI419391 | USDA | Greece |  | <sup>d</sup> |
| *M. prostrata* | PI577445 | USDA | Italy |  | 16<sup>b</sup> |
| *M. rhodopea* | SA43026 | SARDI | NA |  | 16<sup>a</sup> |
| *M. rhodopea* | W619154 | USDA | Bulgaria |  | 16<sup>a</sup> |
| *M. rugosa* | PI487386 | USDA | Tunisia |  | <sup>d</sup> |
| *M. sativa*<sup>c</sup> | PI577567 | USDA | Italy |  | 16<sup>a</sup> |
| *M. scutellata* | PI505433 | USDA | Spain |  | 16<sup>a</sup> |
| *M. suffruticosa* | PI516913 | USDA | Morocco |  | <sup>d</sup> |
| *M. suffruticosa* | W64952 | USDA | Spain |  | 16<sup>a</sup> |
| *M. truncatula* | W64996 | USDA | Greece |  | <sup>d</sup> |

<sup>a</sup>, flow cytometry; <sup>b</sup>, chromosome counts; <sup>c</sup>, five individuals used; <sup>d</sup>, samples confirmed as diploid by inspection of the phased samtools files (sam files) for all eight genes using Tablet (Milne et al., 2013, 2010).

### 2.2 Sequence preparation

We extracted genomic DNA with a custom CTAB DNA Extraction Protocol and constructed a genotyping-by-sequencing (GBS) library following the library preparation protocol of Elshire et al. (2011) with minor modifications as described by Annicchiarico et al. (2017). In brief, GBS library was prepared using the frequent cutter ApeKI (R0643L; NEB) restriction enzyme. Sets of 8-bp barcoded adapters were ligated to restriction fragments for multiplex sequencing. The QIAquick PCR purification kit (28104; QIAGEN) was used to purify equal volumes of the pooled ligated products previous to the final PCR amplification step with the Kapa Library Amplification Readymix (Kapa Biosystems KK2611. Sequences were obtained at the Genomic Core Facility of the UT Southwestern Medical Center (Dallas, TX) with an Illumina HiSeq 2500 system that generated 100-bp single-end reads. This protocol was chosen based on comparisons made among a few protocols and different enzymes, including the two-enzyme protocol by Poland et al. (2012) and the 2b-RAD protocol by Wang et al. (2012). The decision was made based on the number of sites genotyped that were shared among representative individuals (Annicchiarico et al., 2017).

Raw single-end sequence reads were trimmed of adapter sequence and filtered with a minimum quality score of 20 using trimmomatic (Bolger et al., 2014). Assembly was then performed using ipyrad v. 0.7.19 (http://ipyrad.readthedocs.io/), a toolbox for reproducible assembly and analysis of RADseq type genomic data sets based on the pyRAD pipeline (Eaton, 2014). Assembly consisted of seven sequential steps, with parameters based on those recommended for single-end GBS data in the ipyrad documentation. We used the *de novo* + reference method, with the *M. truncatula* genome sequence (Mt4.0, http://www.medicagohapmap.org) as a reference. Briefly, the steps of the ipyrad pipeline are described as follows: In step 1, sequences were demultiplexed according to barcode sequences. In step 2, low quality reads and Illumina adapters were filtered out. Step 3 removed amplification duplicates and then clustered reads within each sample according to a clustering threshold. This step tries to identify all the reads that map to the same locus within each sample. As we used the *de novo* + reference method, the *M. truncatula* reference was used to identify homology, and then the remaining unmatched sequences were clustered with the standard *de novo* ipyrad pipeline. Because phylogenetic results are known to be sensitive to the similarity threshold employed in step 3 for within-sample and step 6 (see below) for across-sample sequence clustering (Fernández-Mazuecos et al., 2017; Leaché et al., 2015; Shafer et al., 2017; Takahashi et al., 2014), five assemblies of GBS loci were generated using a range of clustering thresholds (clust parameter) from c=0.75 to c=0.95 (Table 2). Step 4 jointly estimated the error rate and heterozygosity to differentiate “good” reads from sequencing errors. Step 5 called the consensus of sequences within each cluster. Step 6 clustered consensus sequences across samples. Step 7 filtered the data and wrote output files. In step 7 we applied filters for the maximum number of indels *per* locus (8), max heterozygosity *per* locus (50% of samples) and max number of SNPs *per* locus (20). To evaluate the effect of missing data on network inference, for each assembly we generated datasets with two alternative values for the minimum number of samples *per* locus (“minimum taxon coverage” -min- parameter, 4 and 10). The effect of locus variation on networks was tested by generating datasets with two alternative values for the minimum number of parsimony-informative sites (PIS parameter, 4 and 10). We saved the data in the ipyrad format (*.loci) that was later on transformed in individual alignment files *per* locus in the phylip format using a custom R script. We obtained 20 RADseq datasets under different combinations of assembly parameters and filters described above (Table 2).

**Table 2.** Characteristics of RADseq datasets generated in ipyrad and used for gene tree and network inference.

| Dataset | Clustering threshold | Minimum taxon coverage | Minimum PIS *per* loci | Number of loci | Concatenated length (bp) | Missing data (%) |
| --- | --- | --- | --- | --- | --- | --- |
| clust75.min4.PIS4 | 0.75 | 4 | 4 | 3194 | 283,753 | 49.7 |
| clust75.min4.PIS10 | 0.75 | 4 | 10 | 346 | 29,974 | 56.6 |
| clust75.min10.PIS4 | 0.75 | 10 | 4 | 1756 | 153,733 | 34.4 |
| clust75.min10.PIS10 | 0.75 | 10 | 10 | 140 | 12,274 | 41.5 |
| clust80.min4.PIS4 | 0.80 | 4 | 4 | 3364 | 299,468 | 49.5 |
| clust80.min4.PIS10 | 0.80 | 4 | 10 | 317 | 28,744 | 55.3 |
| clust80.min10.PIS4 | 0.80 | 10 | 4 | 1856 | 162,504 | 34.0 |
| clust80.min10.PIS10 | 0.80 | 10 | 10 | 134 | 11,846 | 40.4 |
| clust85.min4.PIS4 | 0.85 | 4 | 4 | 3405 | 303,272 | 48.8 |
| clust85.min4.PIS10 | 0.85 | 4 | 10 | 205 | 18,789 | 51.8 |
| clust85.min10.PIS4 | 0.85 | 10 | 4 | 1910 | 167,477 | 33.8 |
| clust85.min10.PIS10 | 0.85 | 10 | 10 | 98 | 8,711 | 38.4 |
| clust90.min4.PIS4 | 0.90 | 4 | 4 | 3036 | 271,409 | 48.0 |
| clust90.min4.PIS10 | 0.90 | 4 | 10 | 81 | 7,451 | 45.5 |
| clust90.min10.PIS4 | 0.90 | 10 | 4 | 1729 | 151,806 | 33.3 |
| clust90.min10.PIS10 | 0.90 | 10 | 10 | 40 | 3,616 | 33.2 |
| clust95.min4.PIS4 | 0.95 | 4 | 4 | 1506 | 135,056 | 46.1 |
| clust95.min4.PIS10 | 0.95 | 4 | 10 | 5 | 455 | 24.2 |
| clust95.min10.PIS4 | 0.95 | 10 | 4 | 912 | 80,449 | 32.7 |
| clust95.min10.PIS10 | 0.95 | 10 | 10 | 4 | 367 | 16.2 |

### 2.3 Network inference

We analyzed RADseq datasets alignments with PhyloNet (Than et al., 2008; Wen et al., 2017). Within PhyloNet we applied the method that infers species networks from gene trees using maximum pseudo-likelihood (InferNetwork_MPL command; Yu and Nakhleh, 2015) which is computationally fast. First, we analyzed separate sets of genes from each of the 20 RADseq datasets with RaxML v.7.2.8 (Stamatakis, 2006) using the GTRCAT substitution model and using *M. truncatula* as outgroup. We sampled several individuals/alleles for some species (see Table 1) that were mapped to single taxa with the -a parameter. Ten optimal networks were returned with the -n parameter. We chose 5 maximum allowed number of reticulation events. Remaining parameter values were set as default.

### 2.4 Network distances

To investigate dissimilarities between evolutionary networks computed with alternative RADseq datasets we used multidimensional scaling. We first calculated a matrix of distances among networks computed with the topological dissimilarity measure of Nakhleh (2010) (normalized to get values within [0, 1]), which is implemented in PhyloNet. Then we applied a Principal Coordinate Analysis (PCoA) to transform the distance matrix into a set of coordinates that were plotted to display network distances. We performed the PCoA using all pairwise distances between every network returned by the PhyloNet analyses.

## 3. Results

### 3.1 Sequence capture and RADseq data

Among the 20 RADseq datasets the number of loci ranked from 4 (clust95.min10.PIS10) to 3,405 (clust85.min4.PIS4), concatenated length (bp) ranged from 367 (clust95.min10.PIS10) to 303,272 (clust85.min4.PIS4) and missing data (%) ranged from 16.2 (clust95.min10.PIS10) to 56.6 (clust75.min4.PIS10).

### 3.2 Phylogenetic networks

Best networks (networks with highest likelihood scores) for each of the 20 RADseq datasets showed marked differences (Fig. 1). A hybrid origin was recovered for all species (excluding the outgroup species, *M. truncatula*) at least in one of the 20 best species networks (Table 3): *M. carstiensis* (observed as hybrid in 18 networks), *M. cretacea* (in 16 networks), *M. rhodopea* (in 10 networks), *M. marina* (in 6 networks), *M. rugosa* (in 4 networks), *M. scutellata* (in 4 networks), *M. daghestenica* (in 4 networks), *M. suffruticosa* (in 4 networks), *M. prostrata* (in 2 networks), *M. hybrida* (in 2 networks) and *M. sativa* (in 2 networks).

**Table 3.** Positive hybridization signal detected for each taxon in each network. Only strict hybridization signal is considered, i.e. a taxa nested within a hybrid clade but represented with a single branch is not considered of hybrid origin.

| Dataset | *M. carstiensis* | *M. cretacea* | *M. daghestanica* | *M. hybrida* | *M. marina* | *M. prostrata* | *M. rhodopea* | *M. rugosa* | *M. sativa* | *M. scutellata* | *M. suffruticosa* | *M. truncatula* |
| --- | --- | --- | --- | --- | --- | --- | --- | --- | --- | --- | --- | --- |
| clust75.min4.PIS4 | ✓ | ✓ |  |  |  |  |  | ✓ |  |  |  |  |
| clust75.min4.PIS10 | ✓ |  |  |  |  |  | ✓ |  |  | ✓ |  |  |
| clust75.min10.PIS4 | ✓ | ✓ |  |  |  |  | ✓ | ✓ |  |  |  |  |
| clust75.min10.PIS10 | ✓ | ✓ | ✓ |  | ✓ |  | ✓ |  |  |  |  |  |
| clust80.min4.PIS4 | ✓ | ✓ |  |  |  |  |  |  |  | ✓ |  |  |
| clust80.min4.PIS10 | ✓ |  |  |  |  |  | ✓ |  |  | ✓ |  |  |
| clust80.min10.PIS4 | ✓ | ✓ |  |  |  |  | ✓ |  |  |  |  |  |
| clust80.min10.PIS10 | ✓ | ✓ |  |  |  | ✓ | ✓ |  |  |  | ✓ |  |
| clust85.min4.PIS4 | ✓ | ✓ |  |  |  |  |  |  |  |  | ✓ |  |
| clust85.min4.PIS10 | ✓ |  | ✓ |  | ✓ | ✓ |  |  |  |  |  |  |
| clust85.min10.PIS4 | ✓ | ✓ |  |  |  |  |  |  |  |  |  |  |
| clust85.min10.PIS10 |  | ✓ |  | ✓ | ✓ |  |  |  | ✓ |  |  |  |
| clust90.min4.PIS4 | ✓ | ✓ |  |  |  |  |  |  |  |  | ✓ |  |
| clust90.min4.PIS10 | ✓ | ✓ | ✓ |  | ✓ |  |  |  |  |  |  |  |
| clust90.min10.PIS4 | ✓ | ✓ |  |  | ✓ |  | ✓ |  |  |  |  |  |
| clust90.min10.PIS10 | ✓ |  |  |  |  |  | ✓ |  |  |  | ✓ |  |
| clust95.min4.PIS4 | ✓ | ✓ |  |  | ✓ |  |  | ✓ |  |  |  |  |
| clust95.min4.PIS10 | ✓ | ✓ |  | ✓ |  |  | ✓ |  |  |  |  |  |
| clust95.min10.PIS4 | ✓ | ✓ |  |  |  |  |  | ✓ | ✓ |  |  |  |
| clust95.min10.PIS10 |  | ✓ | ✓ |  |  |  | ✓ |  |  | ✓ |  |  |

**Fig. 1.**
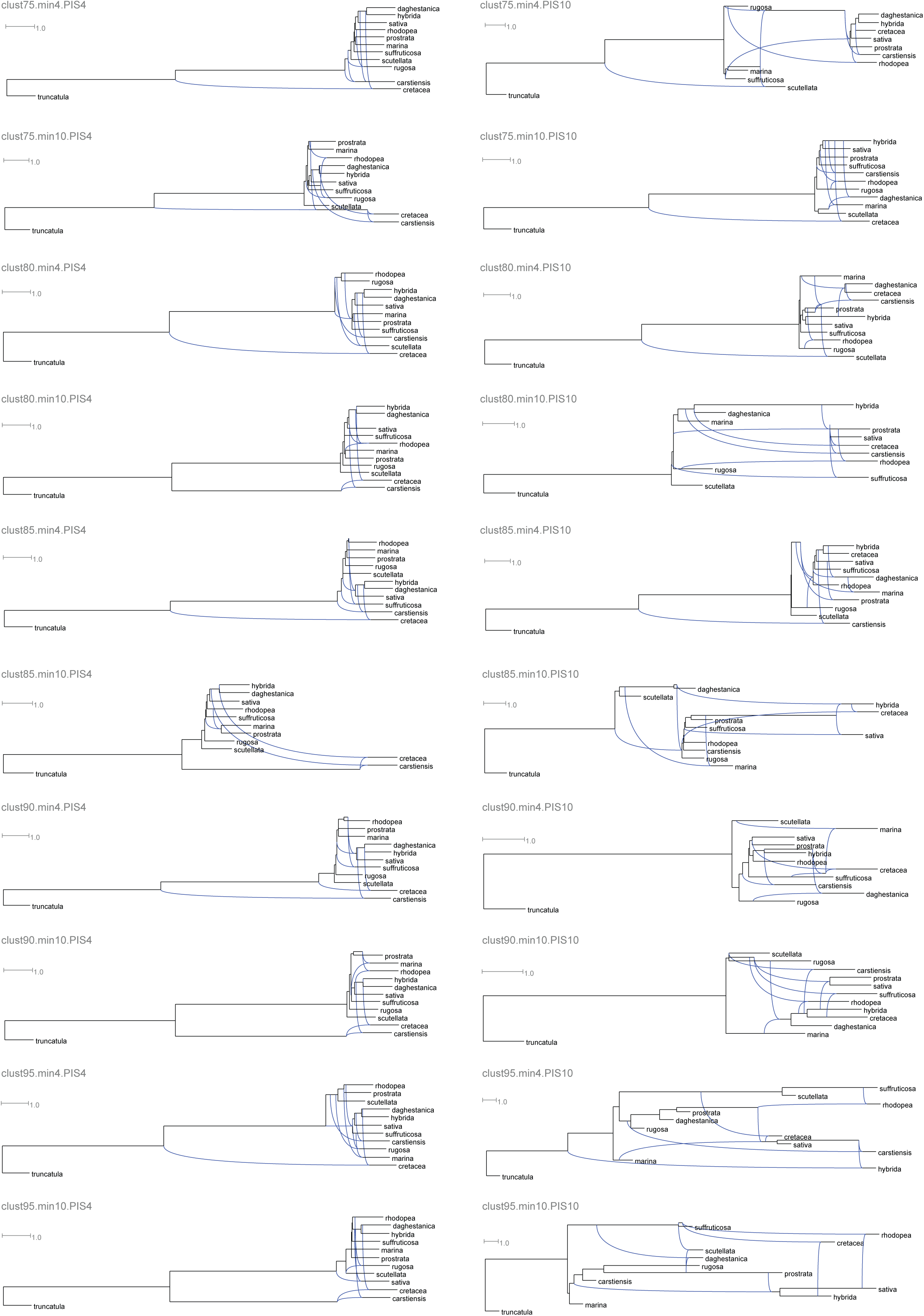
Best networks (networks with highest likelihood scores) for each of the 20 RADseq datasets.

### 3.3 Network distances

The RADseq datasets that retrieved the highest distances from the “core” set of networks were those that were computed with datasets filtered to contain only the most variable loci (PIS10 filter, see Fig. 2). In general, these datasets contained a low number of loci and short concatenated sequence lengths. The PCoA did not show a marked effect of the filter on the minimum number of samples *per* locus (min filter) or the use alternative clustering thresholds (clust parameter).

**Fig. 2.**
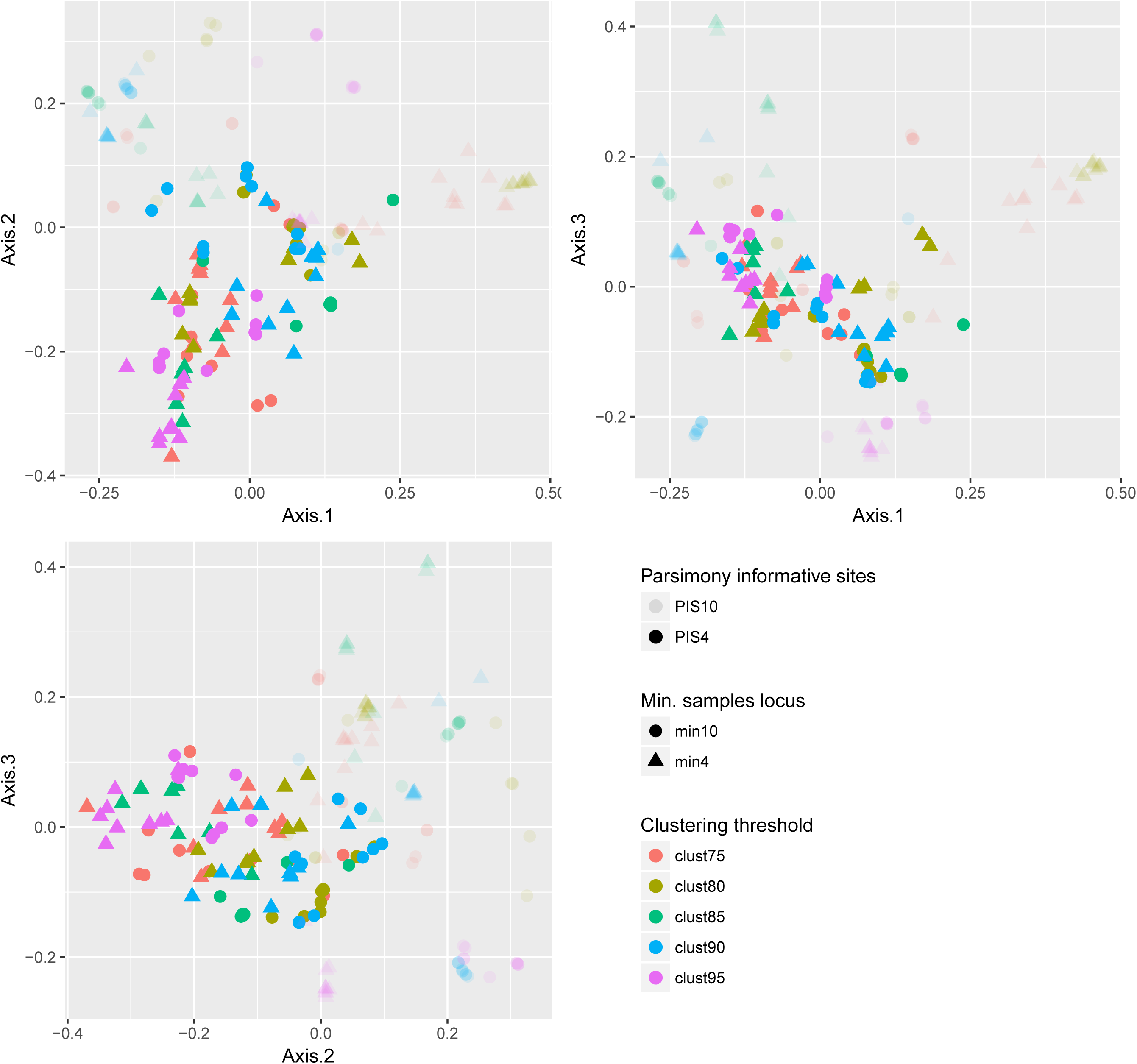
PCoA showing pairwise network distances calculated with the topological dissimilarity measure of Nakhleh (2010). The figure show pairwise distances between the 10 best networks of each of the 20 RADseq datasets. Transparency represents the filter on parsimony informative sites, shapes represent the filter on min. samples locus, and colors represent the clustering threshold used to generate the RADseq dataset.

**Fig. 3.**
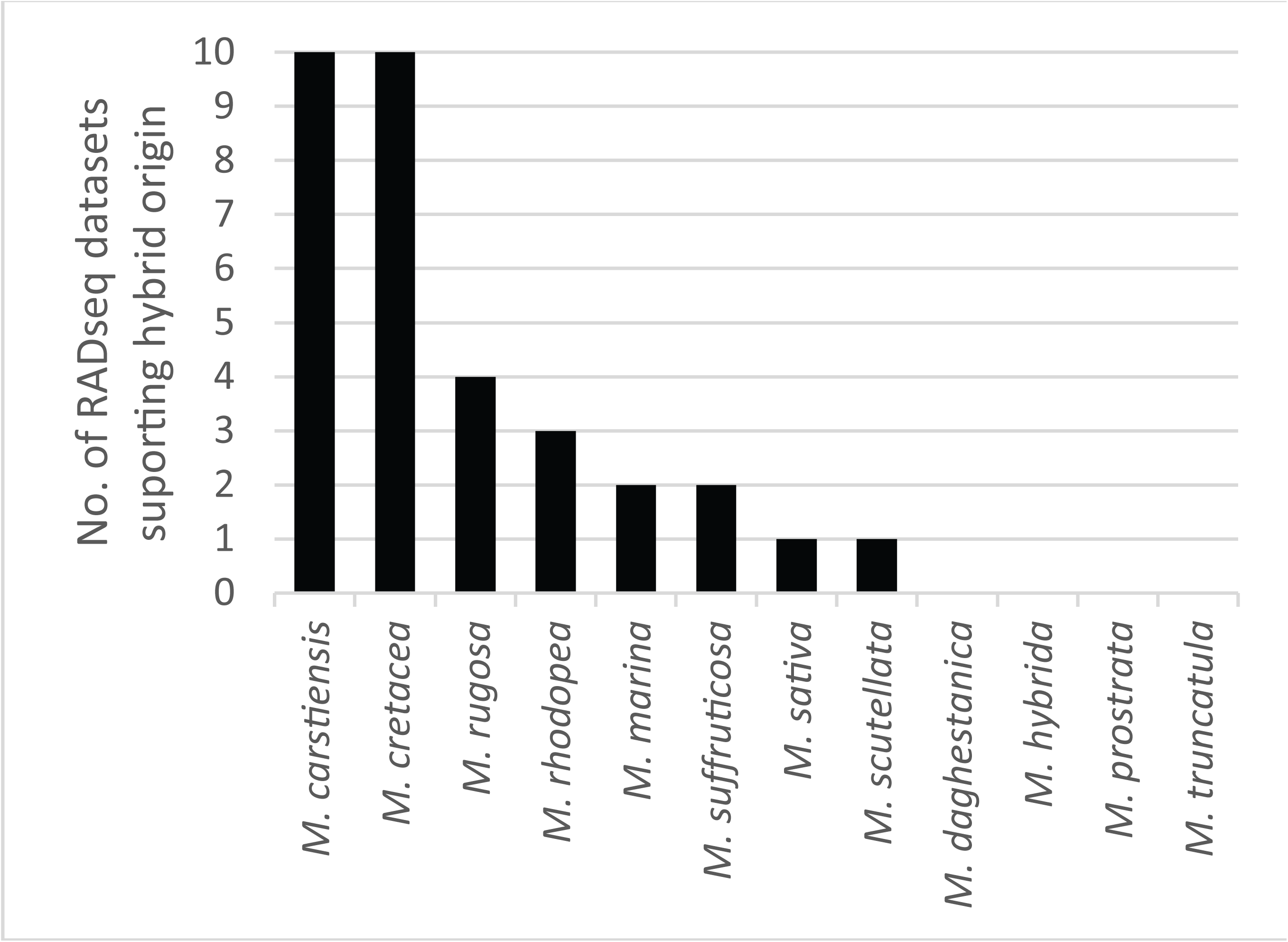
Bar chart representing number of PIS4 RADseq datasets supporting a hybrid origin for the *Medicago* species analyzed in this study.

After excluding datasets with the PIS10 filter, a hybrid origin was recovered for eight species at least in one of the remaining 10 best species networks: *M. carstiensis* (observed as hybrid in all 10 networks), *M. cretacea* (in all 10 networks), *M. rugosa* (in 4 networks), *M. rhodopea* (in 3 networks), *M. marina* (in 2 networks), *M. suffruticosa* (in 2 network), *M. scutellata* (in 1 network) and *M. sativa* (in 1 network).

## 4. Discussion

Our empirical comparison among networks computed from the RADseq datasets reveal some general patterns in how assembly parameters and filters influence complex evolutionary network reconstructions from gene trees. Our study shows that RADseq datasets with a low number of loci retrieve the most atypical network topologies, regardless the high phylogenetic information contained in the loci. The RADseq networks that were the closest to the core set of networks were those that assembled the highest number of loci with very little impact on the clustering threshold or the minimum number of samples *per* locus and therefore with very little impact on the level of missing data. In general RADseq datasets showed low robustness (different best network topologies) under variation of the assembly parameters and filters. But, after excluding the most divergent networks, all remaining analyses supported a hybrid origin for two species: *M. carstiensis* and *M. cretacea*.

Recently Fernández-Mazuecos et al. (2017) showed a high robustness of coalescent approaches for RADseq-based species trees reconstructions. Contrastingly, here we observed variation among network topologies under variations of the assembly parameters and filters underlining the importance of RADseq data preparation on the final results. RADseq is a particularly appealing technique for systematics because of their potential for detecting both current and historical hybridization (Escudero et al., 2014; Twyford and Ennos, 2012) and because they are easily applied with no previous genomic information and reduced lab costs. The pseudo-likelihood method of Yu and Nakhleh (2015) is also attractive because it does not require heavy computational resources. Nevertheless its use on RADseq data may produce misleading results without a proper evaluation of the optimal assembly parameters and filters. In phylogenetic analysis with RADseq, it is particularly challenging to establish general criteria for determining the assembly parameters that maximize the number of orthologous RAD sequences between samples and filtering parameters that retain loci with the optimal level of missing data or phylogenetic information. It has been suggested that low phylogenetic resolution of loci may constrain the identification of hybrids because poorly resolved gene trees, constructed from markers with limited sequence divergence between species, are likely to be uninformative in tracing the reticulate history of species (Linder and Rieseberg, 2004; Twyford and Ennos, 2012). In contrast our study suggests that high loci number increases the power for network inference from RADseq-gene trees despite the low phylogenetic information contained within each individual locus. Additionally, selectively choosing the most variable RADseq dataset may be detrimental as these loci may introduce potential biases typical of hypervariable regions of the genome. Indeed, the most variable regions could be those retaining ancestral polymorphisms or those representing regions of introgressed DNA (Eaton and Ree, 2013).

In recent years RADseq has been applied for the evolutionary reconstruction of complex taxonomic groups (Eaton and Ree, 2013; Escudero et al., 2014; Fernández-Mazuecos et al., 2017; Hipp et al., 2014; Wagner et al., 2013). Most previous studies using RADseq data relied on a “backbone tree” and placed a limited number of hybridization events upon it. Nevertheless it is known that this approach could provide an incorrect representation of the evolutionary history of the species under complex reticulate evolution with multiple hybridization events (Huson et al., 2010; Yu et al., 2011). New tools for evolutionary network reconstructions (Solís-Lemus and Ané, 2016; Wen et al., 2016; Wen and Nakhleh, 2016; Yu et al., 2014, 2013; Yu and Nakhleh, 2015; Zhang et al., 2018; Zhu et al., 2017) now offer the opportunity to study reticulate evolution including cases with multiple hybridization events and with no previous information on the “backbone tree” or where such main tree is potentially non-existent. Development of such evolutionary network models are now in full swing and should become standard methods for phylogenetic inference under incomplete lineage sorting (ILS) and hybridization. Despite these remarkable methodological advances, in the most complex cases computational limitations reduce the set of methods to those using maximum pseudo-likelihood inference of networks from gene trees (Solís-Lemus and Ané, 2016; Yu and Nakhleh, 2015). These methods have a great potential but there is no information in the literature about the adequacy of the commonly used RADseq datasets for the estimation of evolutionary networks using these type of analyses were a previous computation of gene trees is required. In general using RADseq for gene tree reconstruction poses a number of potential problems: the orthology relationships among sequences are unknown, mutations on restriction sites is expected to yield missing data that increases with evolutionary time and the genetic linkage relationships among loci are unknown (see Rubin et al., 2012). Additionally, given the short length of sequences, the phylogenetic information of each locus is very scarce and recombination detection is not straightforward.

Despite the varied result obtained with different RADseq datasets, general patterns emerged regarding the identification of hybrid species which were more evident when the PIS10 datasets were excluded. A hybrid origin was retrieved by all remaining PIS4 datasets for *M. carstiensis* and *M. cretacea*. This signal was clearly stronger than the hybridization signal detected for the remaining species (hybrid origin detected in ≤ 4 datasets for *M. rugosa, M. rhodopea, M. suffruticosa, M. marina, M. scutellata* and *M. sativa*). *M. carstiensis* is the only *Medicago* species exclusively with rhizomes (which are found only sporadically in a few other species, especially *M. sativa*). Phylogenetic relationships around *M. carstiensis* has been enigmatic as it forms a monospecific section (Carstiensae, Small, 2011) and previous phylogenetic studies did not provide well-supported information on the relationships of this species (Maureira-Butler et al., 2008; Small, 2011). It has been speculated that *M. carstiensis* is a relic species that is ancestral to the much more widespread *M. orbicularis* (Bennett et al., 2006). But its particular characteristics could be also explained by speciation after disruptive selection on hybrids (Seehausen, 2004). Regarding *M. cretacea*, Urban (1873) (the first to prepare a comprehensive analysis of the genus *Medicago*) already considered that this species had controversial affinities. Lesins and Lesins (Lesins and Lesins, 1979), in the second comprehensive systematic analysis of *Medicago*, already included *M. cretacea* in the monotypic section *Cretaceae*. Later on, analyses by Bena (2001) and Maureira-Butler et al. (2008) showed alternative inconsistent phylogenetic relationships for *M. cretacea*. These contentious taxonomic and phylogenetic placement of *M. carstiensis* and *M. cretacea* observed in previous studies are consistent with hybridisation.

## 5. Conclusions

Here we inferred a hybrid origin for *M. carstiensis* and *M. cretacea* using RADseq data and a maximum pseudo-likelihood approach for network inference from gene trees. We observed that loci number had an important impact on network reconstruction from RADseq-gene trees, whereas the clustering threshold used in the data assembly or a filter on taxon coverage had a lower impact on network inference. Future research on methods that explore the parameter space for optimal assembly parameters and filters may be required to obtain a clear phylogenetic picture of all diploid perennial *Medicago* species and to consider these approaches sufficiently robust for their standard use in the phylogenetics community.

## Acknowledgements

The authors thank Luay Nakhleh and Jiafan Zhu for their assistance with PhyloNet analyses and Filipe de Sousa for providing plant material.

